# Phthalate monoesters affect membrane fluidity and cell-cell contacts in endometrial stromal cell lines

**DOI:** 10.1101/2024.06.17.599271

**Authors:** Darja Lavogina, Keiu Kask, Sergei Kopanchuk, Nadja Visser, Mary Laws, Jodi A. Flaws, Theodora Kunovac Kallak, Matts Olovsson, Pauliina Damdimopoulou, Andres Salumets

## Abstract

Phthalate monoesters have been identified as endocrine disruptors in a variety of models, yet understanding of their exact mechanisms of action and molecular targets in cells remains incomplete. Here, we set to determine whether epidemiologically relevant mono(2-ethyl-5-hydroxyhexyl) phthalate (MEHHP) can affect biological processes by altering cell plasma membrane fluidity or formation of cell-cell contacts. As a model system, we chose endometrial stromal cell lines, one of which was previously used in a transcriptomic study with MEHHP or MEHHP-containing mixtures. A short-term exposure (1 h) of membrane preparations to endocrine disruptors was sufficient to induce changes in membrane fluidity/rigidity, whereas different mixtures showed different effects at various depths of the bilayer. A longer exposure (96 h) affected the ability of cells to form spheroids and highlighted issues with membrane integrity in loosely assembled spheroids. Finally, in spheroids assembled from T-HESC cells, MEHHP interfered with the formation of tight junctions as indicated by the immunostaining of *zonula occludens* 1 protein. Overall, this study emphasized the need to consider plasma membrane, membrane-bound organelles, and secretory vesicles as possible biological targets of endocrine disruptors and offered an explanation for a multitude of endocrine disruptor roles documented earlier.

## Introduction

Endocrine disruptors are individual chemicals or chemical mixtures that mimic hormones (especially steroid hormones) and thus affect endocrine signalling in organisms leading to adverse health outcomes (1). This term covers a broad range of chemical scaffolds featuring different ADME-Tox properties (2). One of the well-known classes of endocrine disruptors is represented by phthalic acid esters which are widely used as plastic constituents in a variety of industrial products (3) but also as additives and fixatives in cosmetics (4). In organisms, phthalates are relatively quickly metabolized to phthalic acid monoesters (5) which have been shown to be endocrine disruptors in a wide choice of model systems, especially those related to reproductive health. Less data are available on mixtures of phthalates, although organisms are usually exposed to a variety of different endocrine disruptors during the lifetime (6).

The studies in simplified models have so far not pinpointed a single target or a group of biological targets affected by phthalates (7, 8). Given their physicochemical properties (*e*.*g*., low molecular weight, semi-volatility, limited solubility, variable hydrophobicity, incorporation of a small aromatic system and side-chain functional groups that can be oxidation-prone) (9, 10), it is not surprising that phthalates can bind multiple targets within a physiologically meaningful range of affinities, thus obscuring the interpretation of omics data. Our recent publications focusing on the effects of epidemiologically relevant phthalate mixtures at the transcriptome level reported cell model-dependent mode of action in the case of human and mouse ovaries, indicating the involvement of phthalates in processes related to cell adhesion and cytoskeleton functioning (11, 12). This observation motivates further studies of phthalate effects on cell adhesion and cytoskeleton changes in endometrial cells, where extensive morphological alterations need to take place every month as a part of the menstrual cycle (13).

In principle, the range of putative targets of endocrine disruptors in general and phthalates in particular are not limited to receptors, enzymes, and transcription factors. Endocrine disruptors can also affect biological membranes by intercalating into the bilayer (14) and altering membrane fluidity and formation of lipid rafts (15, 16). Previous studies on the membrane-related effects of endocrine disruptors and similar compounds are available mainly for bisphenol A (17), azelaic acid esters (18) and some steroid derivatives (15), whereas phthalate monoesters have only been examined in the context of bacterial membranes (19). Still, given the structural similarities of phthalate monoesters and major components of the mammalian plasma membrane (phospholipids and cholesterol), it is likely that phthalates can be incorporated into the bilayer structure and affect the membrane properties. Importantly, the composition and fluidity of biological membranes (*i*.*e*., not only plasma membrane, but also mitochondrial membrane) are connected to a multitude of outcomes on the cellular level – such as cell surface receptor-initiated signalling (20), steroidogenesis (21) and other metabolic processes (22), cell cycle (23), secretion (24) and cell-cell signalling (25), etc. Thus, we hypothesize that effects of phthalates on membrane fluidity explain the reported involvement of these chemicals in a variety of biological processes, including those directly related to cell adhesion.

Here, we set out to determine whether phthalate monoesters can affect biological systems by inducing changes in membrane fluidity and cell-cell contacts. For our research, we chose mono(2-ethyl-5-hydroxyhexyl) phthalate and phthalate mixtures that we used in two previous studies exploring short-term exposure effects at the transcriptome level (11, 12). In the current study, we utilized both short-term and long-term exposures, which were adjusted according to the model system of choice (biological membranes *versus* cell spheroids). Due to the requirement for a large amount of cells in membrane fluidity assays, we conducted the experiments in endometrial stromal cell lines rather than primary cells.

## 2. Materials and methods

## 2.1. Apparatuses and reagents

The 1000× stock solution of mono(2-ethyl-5-hydroxyhexyl) phthalate (MEHHP) and mixtures of phthalate monoesters in DMSO (Sigma-Aldrich, USA) were prepared and validated as previously reported (11). The epidemiological mixture (EPIDEM) and the equimolar mixture (EQUI) contained MEHHP, mono(ethylhexyl) phthalate (MEHP), mono(2-ethyl-5-oxyhexyl) phthalate (MEOHP), mono(5-carboxy-2ethylpentyl) phthalate (MECPP), monoisobutyl phthalate (MiBP), monobenzyl phthalate (MBzP), mono(3-carboxypropyl) phthalate (MCPP), monoethyl phthalate (MEP), and monobutyl phthalate (MBP). The weighted quantile sum-based mixture (WQS) contained MEHHP, MiBP, MBzP and MEP (see Supplementary Table S1 for details). Mixtures were distributed into 10Lμ□ aliquots and stored at -20□°C.

The telomerase reverse transcriptase-immortalized human endometrial stromal cell line T-HESC was purchased from the American Type Culture Collection (ATCC CRL-4003). The immortalized human endometrial stromal cell line St-T1b was a kind gift from Dr. Martin Götte (University of Münster, Germany) (26). The cell lines were cultured at 37 °C in a 5% CO_2_ humidified atmosphere incubator (Sanyo, Japan). As the growth medium, phenol red-containing Dulbecco’s Modified Eagle’s medium (DMEM) / Ham’s F12 medium (1:1; Corning, USA) supplemented with 10% fetal bovine serum (FBS; Capricorn, Germany), insulin (Sigma-Aldrich, USA) and a mixture of penicillin, streptomycin, and amphotericin B (Capricorn, Germany) was utilized. Prior to membrane preparation, the cells were scaled up at the usual growth conditions using the phenol-red free DMEM/Ham’s F12 (Sigma-Aldrich, Germany).

For preparation of membrane homogenates, the following lysis buffer components were used: phosphate buffered saline (PBS) supplemented with Ca^2+^ and Mg^2+^ (Corning, USA), phenylmethylsulfonyl fluoride from AppliChem (Germany), and cOmplete™ protease inhibitor cocktail supplemented with EDTA from Roche (Switzerland). Homogenization was conducted using the ultra-turrax (Cole-Palmer 3771, Germany); a refrigerated high-speed centrifuge (Sigma 3-30 KS, Germany) together with rotor 12154-H was used for pelleting and washes. The membrane fluidity measurements were carried out in 384-well black round-bottom low volume plates (Corning 4514, USA) using phosphate buffered saline (PBS) supplemented with Ca^2+^ and Mg^2+^ (Corning, USA) as the working buffer and 1,6-diphenyl-1,3,5-hexatriene (DPH) and N,N,N-trimethyl-4-(6-phenyl-1,3,5-hexatrien-1-yl)phenylammonium p-toluenesulfonate (TMA-DPH) as the probes (both from Sigma, USA). As the positive controls, sodium dodecyl sulphate (SDS; Sigma, USA) and Triton X-100 (Ferak, Germany) were utilized. The anisotropy was measured at 30 °C using PHERAstar multi-mode reader (BMG Labtech, Germany) with the filter block FP608B (excitation at 360 nm, emission at 430(10) nm, 200 flashes).

The spheroid formation was conducted on 96-well black ultra-low attachment (ULA) spheroid microplates with clear round bottom (Corning 4515, USA). For staining of cell population with weakened membrane integrity, propidium iodide (PI; Acros Organics, Switzerland) was used. Live imaging of spheroids was performed in an automated focusing mode with Cytation 5 multi-mode reader at 37 °C in a 5% CO_2_ humidified atmosphere, using 4× air objective (1.613 μm/pixel) and bright-field (LED intensity 5, integration time 100, detector gain 3) or RFP channel (LED intensity 5, integration time 350, detector gain 15).

For fixation and permeabilization of spheroids prior to immunostaining, 4% paraformaldehyde (PFA; Thermo Scientific, USA) and Triton X-100 (Naxo, Estonia) in PBS (Thermo Scientific, USA) were utilized, respectively. The immunostaining of tight junction proteins was conducted using rabbit monoclonal antibody against human *Zonula Occludens* 1 protein (ZO-1; Abcam ab221547, UK), rabbit monoclonal antibody against human Occludin (Abcam ab222691, UK), or rabbit monoclonal antibody against human Claudin 1 (Abcam, ab15098, UK). Goat anti-rabbit antibody conjugated with Alexa Fluor 488 (Thermo Scientific, USA) was used as the secondary antibody and Hoechst 33342 (Molecular Probes, USA) was applied for staining of nuclei. The spheroids were mounted onto the specimen slides with 10 reaction wells (Marienfeld, Germany) using Fluoromount G mounting media (Thermo Scientific, USA).

The imaging of fixed spheroids was conducted with the microscopy setup as described previously (27). Briefly, widefield epifluorescence and TIRF were conducted using an inverted microscope built around a Till iMIC body (Till Photonics/FEI, Germany), equipped with UPlanFLN 10×□air (numerical aperture 0.30) objective lens (Olympus Corp., Japan). The samples were sequentially excited with 405 nm (150 mW) and 488 nm (100 mW) PhoxX laser diodes combined in the SOLE-6 light engine (Omicron-Laserage, Germany). Excitation light was launched into Yanus scan head, which along with a Polytrope galvanometric mirror (Till Photonics/FEI, Germany) was used to position the laser for widefield epifluorescence. Excitation and emission light were spectrally separated using imaging filter cubes with the following parameters: for Hoechst 33342 staining (excited with 405 nm laser), a 390/40 BrightLine exciter filter (Semrock, USA), a zt 405 rdc 2 mm thick beamsplitter and a ET 460/50 emission filter (Chroma Technology Corp, USA); for secondary antibody staining (excited with 488 nm laser), a zt 491rdc 2 mm thick beamsplitter (Chroma Technology Corp, USA) and a 525/45 BrightLine bandpass emission filter (Semrock, USA). The electron-multiplying charge-coupled device Ultra 897 camera (Andor Technology, UK) was mounted to a microscope through a TuCam adapter with 2×□magnification (Andor Technology, UK). The camera was cooled down to□−100 °C with the assistance of a liquid recirculating chiller Oasis 160 (Solid State Cooling Systems, USA).

### 2.2. Membrane fluidity assay in T-HESC and St-T1b membrane homogenates

For a single preparation of membrane homogenates, T-HESC or St-T1b cells from three confluent 10 cm Petri dishes were pooled following the trypsinization. The cells were pelleted by centrifugation (800 rcf, 5 min) at room temperature (RT), washed twice with PBS, pelleted again and kept on ice in the subsequent steps. To the cell pellet, 1.5 mL of ice-cold lysis buffer was added; the cells were resuspended and homogenized mechanically (5× 30 second cycles). The membranes were then pelleted by centrifugation (27,000 rcf, 40 min, 4 °C) and the supernatant was discarded. The pellet was resuspended in ice-cold lysis buffer and homogenized as described above; the centrifugation, resuspension and homogenization steps were then repeated. The obtained membrane homogenate was then stored at -80 °C until further usage.

On the day of membrane fluidity measurement, the membrane homogenates were thawed on ice. The measurement was carried out according to the previously reported (28) adjusted protocol. The total working volume was 20 μL (10 μL of a compound or a mixture of interest or assay buffer in the case of negative control, 5 μL of membrane homogenate and 5 μL of probe solution). The following final total concentrations were used: 16 nM DPH or 40 nM TMA-DPH; 1:1000 dilution of MEHHP stock (21.6 μM), 1:1000 dilution of phthalate mixtures’ stocks (net phthalate concentration of 280 μM for EPIDEM, 330 μM for EQUI, or 220 μM for WQS), 20 mM SDS, or 10 mM Triton X-100. The system was preincubated at 30 °C for 60 min before the measurement. In each experiment, the anisotropy values were adjusted by the well containing membrane homogenates and 16 nM DPH (in the absence of detergents; value set to 250 mP units).

In a single pilot experiment for optimization of workflow (Supplementary Figure S1), 2-fold dilution series of SDS in PBS (final total concentration starting from 20 mM) were also tested according to the same measurement protocol. The anisotropy was measured as mentioned above after 30 min or 60 min preincubation.

### 2.3. Spheroid formation assay in T-HESC and St-T1b cell lines

T-HESC or St-T1b cells were seeded in a growth medium onto the 96-well ULA plate with the density of 4000 to 7500 cells per well (different seeding densities were used for different experiments; N = 5). At the same time, 1:1000 stock dilutions of MEHHP (final concentration of 21.6 μM), phthalate mixtures (net phthalate concentration of 280 μM for EPIDEM, 330 μM for EQUI, or 220 μM for WQS), DMSO (final concentration of 0.1%, negative control) or SDS (final concentration of 350 μM, positive control) in the growth medium were added to the wells. The total working volume was 200 μL per well. The plates were incubated for 48 h at 37 °C in a 5% CO_2_ humidified atmosphere; next, half of the medium was replaced with the fresh one (not supplemented with phenol red but containing the same final concentrations of the compounds as outlined above) and the incubation was continued. The switch to phenol red-free medium was required to minimize the background in the PI imaging in the next step. Following the 95-h incubation post-seeding, a concentrated solution of PI in PBS was added to all wells (final total concentration of 2 μg/mL). The plates were incubated for 1 h at 37 °C in a 5% CO_2_ humidified atmosphere and then imaged.

In a single pilot experiment for optimization of the workflow (Supplementary Figure S2), bright-field imaging (without the preceding PI treatment) was also performed at 48 h and 72 h after seeding of cells onto ULA plates. No treatment with compounds of interest was used (*i*.*e*., only growth medium was added to the cell suspension during the cell seeding and medium exchange steps).

### 2.4. Immunostaining of ZO-1 in fixed T-HESC spheroids

T-HESC cells were seeded in growth medium onto the 96-well ULA plate with the density of 7500 cells per well. At the same time, 1:1000 dilutions of MEHHP (final concentration of 21.6 μM) or DMSO (final concentration of 0.1%, negative control) in the phenol red-containing growth medium were added to the wells; the total working volume was 200 μL per well. The plates were incubated for 48 h at 37 °C in a 5% CO_2_ humidified atmosphere; next, half of the medium was replaced with the fresh one (containing the same final concentrations of the compounds as outlined above) and the incubation was continued. The exposure time was reduced by 24 h compared to the spheroid formation assayas the MEHHP-exposed spheroids were too fragile to handle following the longer exposure.

Following the 72-h incubation post-seeding, the spheroids were fixed in 4% PFA for 30 minutes (min) and washed 2× 5 min with PBS. For permeabilization, the samples were incubated in 0.1% solution of Triton X-100 in PBS for 10 min. For non-specific blocking, 30 min treatment with 10% normal goat serum was used, followed by the overnight incubation with primary antibody against ZO-1 tight junction protein (1:100 dilution) at 4 °C. Subsequently, the spheroids were incubated with goat anti-rabbit Alexa Fluor 488 secondary antibody (1:1000 dilution) for 1 h at RT in the dark. The cell nuclei were counterstained by Hoechst 33342 for 3 min in the dark. Finally, the spheroids were transferred one by one to the wells of the specimen slide wells and mounted using Fluoromount G mounting medium. The mounted spheroids were stored at 4 °C until imaging.

In a single pilot experiment for optimization of the workflow (Supplementary Figure S3), non-treated T-HESC and St-T1b spheroids were generated and fixed according to the same protocol and immunostaining of claudin and occludin (1:500 dilution of primary antibody) was additionally performed.

### 2.5. Data analysis and software

For general data analysis, GraphPad Prism 6 (USA) and Excel 2016 (Microsoft Office 365, USA) were used.

In membrane fluidity assays, the anisotropy values in each independent experiment were normalized separately for DPH and TMA-DPH: the negative control (wells with the corresponding probe and membrane homogenate) signal was set to 100%, and the positive control (wells with the corresponding probe, membrane homogenate and 20 mM SDS) signal was set to 0%. The normalized data were then pooled for all replicates in all independent experiments (N ≥ 3 for each compound in each cell line). The statistical significance of pairwise comparisons for the same probe (negative control *versus* compound-treated membranes from the same cell line) was established using the unpaired two-tailed t-test with Welch’s correction: ^***^ indicates P < 0.001, ^**^ indicates P < 0.01, ^*^ indicates P < 0.05, ns indicates not significant.

In spheroid formation assays, ImageJ software (Fiji package) (29) was used for the image analysis. The quantification was performed with minor modifications to the previously reported protocol (30) due to differences in spheroid morphology in the two cell lines. In the case of T-HESCs, all dark area occupied by the cells was assumed to represent the spheroid surface, whereas in the case of St-T1b, the well-defined border between the spheroid and the surrounding liquid was used. The spheroid contour was denoted manually using the freehand selections tool and the area and circularity of the spheroid were quantified. For the RFP channel images, the average signal intensity of PI per unit of spheroid area was also established. The measured parameters were normalized to the negative control in the given cell line (set to 100%) in each independent experiment (N = 5), and the data for the identically treated spheroids within the same cell line (at least 9 spheroids per compound) were then pooled for all independent experiments. The statistical significance of pairwise comparisons (negative control *versus* compound-treated spheroids from the same cell line) was established using the unpaired two-tailed t-test with Welch’s correction: ^***^ indicates P < 0.001, ^**^ indicates P < 0.01, ^*^ indicates P < 0.05, ns indicates not significant.

The microscopic images of fixed spheroids were captured as 16-bit OME-TIFF multicolor Z-stacks (100 frames, with 1 μm focusing increment), using 100 ms exposure time and EM gain 300. Prior to the imaging, the brightfield channel was used for the determination of the spheroid exact location in the well. The widefield epifluorescence Z-stacks were deconvolved with the EpiDEMIC plugin (31) in an ICY platform (32).

For establishing the MEHHP effect on the ZO-1 staining in spheroids, ImageJ software was used. The average intensity of immunostaining in the secondary antibody channel was quantified in raw non-deconvolved images (a single focal plane with the highest total intensity within the Z-stack) by drawing an oval according to the minimal inner diameter of the spheroid. The measured intensity was normalized to the negative control (set to 100%) in each independent experiment (N = 3), and the data was then pooled for all independent experiments. The statistical significance of pairwise comparisons (negative control *versus* MEHHP-treated spheroids) was established using the unpaired two-tailed t-test with Welch’s correction: ^**^ indicates P < 0.01.

## 3. Results

### 3.1. Phthalate monoester mixtures affect differently fluidity of endometrial stromal cell membranes

We started our study by examining the effects of phthalates on the fluidity of membranes isolated from the endometrial stroma-based immortalized cell lines T-HESC and St-T1b. For that, we utilized probes DPH and TMA-DPH that become fluorescent when incorporated into the membrane bilayer. In previous studies by us (28) and other groups (33, 34), it was shown that the fluorescence polarization/anisotropy value (FA) of these probes is strongly dependent on the viscosity of the membrane. Namely, the latter affects the rotational freedom of the probes and hence their ability to emit light that is polarized in the analogous way as the light used for excitation of the probes. Importantly, the two probes incorporate themselves into the membrane bilayer at different depths, with non-charged DPH residing deeper than the positively charged TMA-DPH that can interact with the negatively charged phospholipid heads on the outer part of the bilayer (34, 35). Due to potentially high non-specific binding and low excitation wavelength, the probes cannot be used in measurements with live cells; therefore, a simplified model system represented by the membrane preparations must be utilized.

Initially, we established the effect of a well-known detergent SDS on the DPH and TMA-DPH FA signal in membrane preparations and confirmed the SDS concentration-dependent decrease in membrane rigidity (Supplementary Figure S1). Consistently with the previously reported data(36) and the higher molecular weight of the probe, in the case of TMA-DPH, the measured absolute anisotropy values were systematically higher than in the case of DPH. Next, we performed membrane fluidity measurements in the presence or absence of MEHHP or MEHHP-containing mixtures (EQUI, EPIDEM, or WQS). The normalized results pooled from three or more independent experiments are shown in Figure 1.

**Figure 1.**
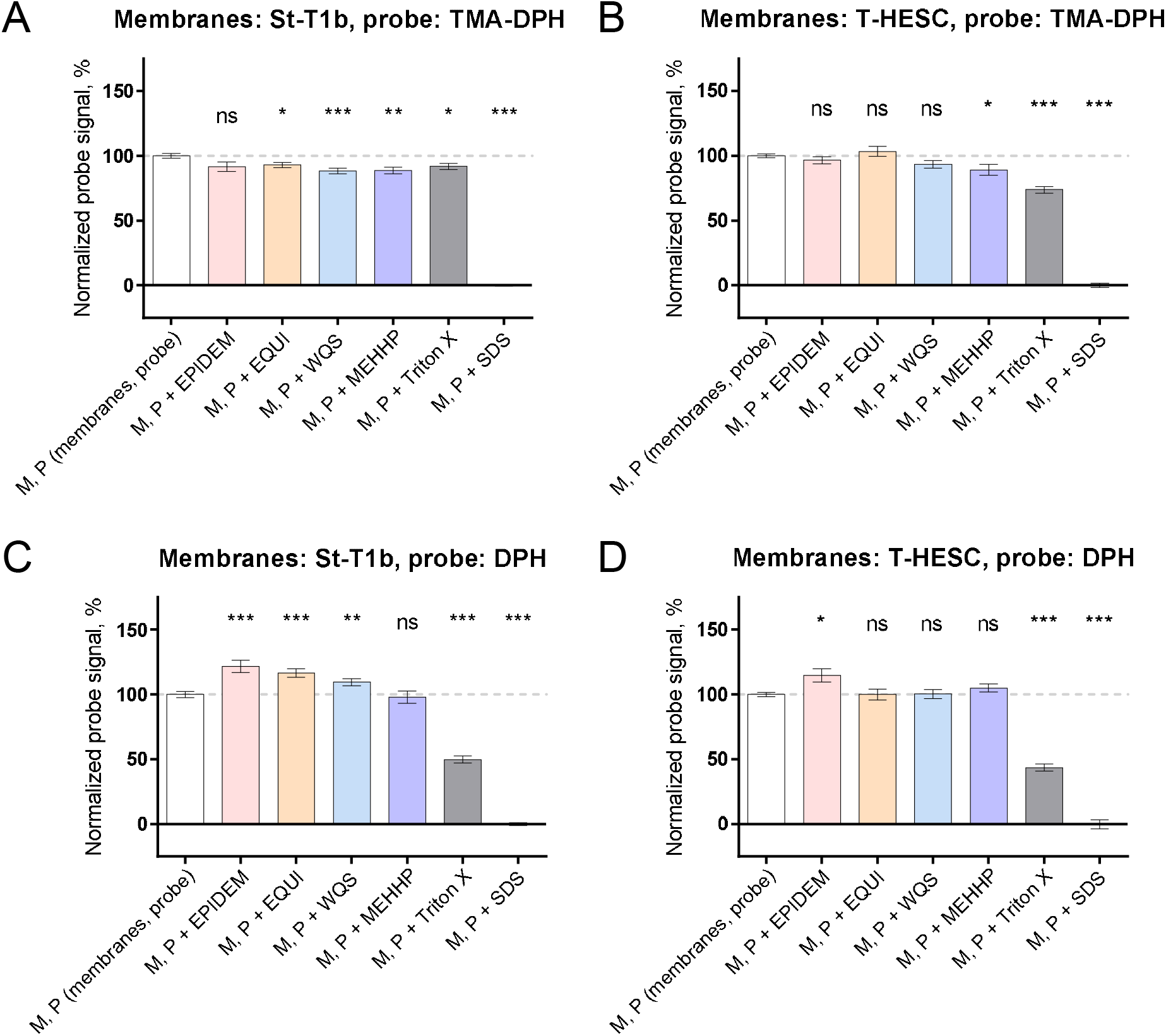
Membrane fluidity assay illustrates changes in fluorescence anisotropy of membrane-intercalating probes in the presence of compounds of interest in endometrial stromal cell membrane preparations. Average normalized anisotropy ± SEM is shown (N ≥ 3 for each compound of interest in each cell line). The compounds of interest are listed below each graph; net concentrations of phthalates: MEHHP – 21.6 μM, EPIDEM – 280 μM, EQUI – 330 μM, WQS – 220 μM. Probes: A-B, TMA-DPH (binds closer to bilayer surface), C-D, DPH (buried deeper inside bilayer). Cell lines: A and C, St-T1b; B and D, T-HESC. The dashed grey line indicates normalized probe signal in the presence of membranes but in the absence of a compound of interest. The asterisks indicate the statistical significance of difference between the normalized probe signal in the absence *vs* presence of the compound of interest; ^***^ indicates P < 0.001, ^**^ indicates P < 0.01, ^*^ indicates P < 0.05, and ns indicates not significant.

Overall, all tested compounds affected membrane fluidity according to measurements with at least one probe in one cell line samples, whereas the differences between the different treatments were more pronounced than the differences between the cell lines. Still, the statistical significance of the compound effect as compared to the non-treated membranes was more readily observed in St-T1b rather than in T-HESC samples. In contrast to classical detergents such as SDS or Triton X that reduced fluorescence anisotropy in the case of both probes (P < 0.05), thus indicating membrane-fluidifying effects at different depths in the bilayer, the effect of phthalates was also depth-dependent. Both EQUI and WQS fluidified the outer part of the membrane (P < 0.05) layer yet rigidified the inner part (P < 0.01) in St-T1b samples; in the case of EPIDEM, only rigidifying effect was observed in the inner part of membranes derived from both cell lines (P < 0.05). MEHHP, on the other hand, showed only fluidifying effect in the outer part of membranes derived from both cell lines (P < 0.05).

The characteristic effects of different treatments in membranes of endometrial stromal cells were visualized after pooling the normalized signal for identically treated membrane preparations from both cell lines in the case of the same probe. The data are presented in Figure 2. The pooling of data highlighted the differences in phthalate monoester treatment as compared to the classical detergents and indicated that while MEHHP and WQS mixture had similar effects, the EQUI mixture was the most different from the other treatments.

**Figure 2.**
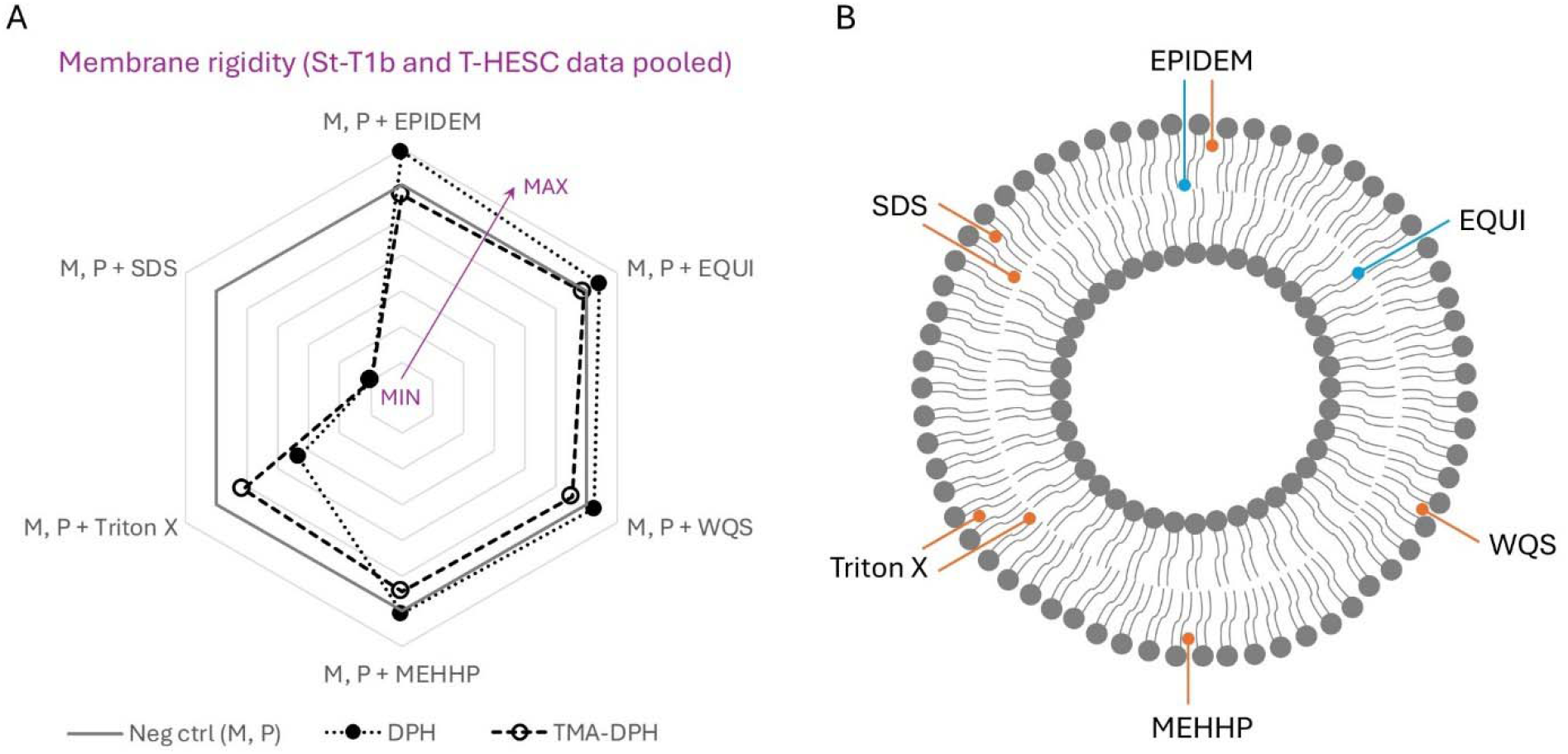
Characteristic fingerprint of different phthalate monoesters in the context of biological membranes. A, Radar map based on membrane fluidity assay data pooled for two cell lines used. The higher is the probe signal, the lower is membrane fluidity and the higher is membrane rigidity. Each data point represents normalized mean probe signal in the case of a given treatment (used mixtures and individual compounds listed on the diagram periphery); the error bars were omitted for the purpose of better readability. M stands for membrane homogenate and P for probe, the probes are listed below the graph. B, Working model of MEHHP and MEHHP-containing mixture effects in the context of biological membranes. The model is based on the data shown in part A. The location of arrow ends indicates different depths in the lipid bilayer (according to the data measured using DPH and TMA-DPH); blue arrows indicate rigidifying effect and orange arrows fluidifying effect.

### 3.2. Phthalate monoesters affect spheroid formation from live endometrial stromal cells

Next, we tested whether the effects of phthalates observed in a simplified cell-free system represented by the membrane preparations are reflected at the level of live cells. As alterations in membrane fluidity or rigidity are known to affect assembly of cells into spheroids (37, 38), we performed spheroid formation assay where the freshly trypsinized suspension of St-T1b or T-HESC cells was mixed with a compound of interest and immediately seeded onto the ULA plate. In a pilot experiment, the compounds of interest were represented by the 0.1% DMSO (negative control) and 350 μM SDS (positive control); the forming spheroids were imaged at 48 h, 72 h, and 96 h post-seeding. In subsequent experiments, in addition to the control treatments, MEHHP and the phthalate mixtures were explored as the compounds of interest; following the 95-h incubation, the spheroids were stained with PI and then imaged using bright-field microscopy (to measure the area and circularity of the formed spheroid) and in parallel with fluorescent microscopy (to measure the intensity of PI staining).

The pilot experiment indicated that the morphology of non-treated spheroids formed by cells of two different cell lines varied remarkably already after 48 h post-seeding and even more upon prolonged incubation (Supplementary Figure S2). The St-T1b cells formed compact spheroids with a well-outlined surface, whereas the T-HESC cells formed less compact and less circular spheroids. Therefore, two different image analysis approaches were used for the St-T1b and the T-HESC spheroids in the subsequent experiments. For St-T1b, the area of the spheroid was contoured along the well-defined borderline (except in the case of SDS treatment where no clear surface was formed), whereas for T-HESC, all dark area occupied by the spheroid-forming cells (including the protrusions) was quantified. For better comparison of independent experiments (N ≥ 3 for each compound in each cell line), the values of parameters measured in each experiment were normalized by the negative control.

The data on the normalized spheroid area, circularity and average intensity of PI staining per area unit are presented in Figure 3; the representative images of live spheroids are shown in Figure 4. Interestingly, the effects of different phthalate monoester mixtures (EQUI, EPIDEM, WQS) and MEHHP alone were similar in the spheroid formation assay. In the case of St-T1b cell line, the area of the well-defined spheroid was slightly yet significantly reduced by EQUI, WQS, and MEHHP (P < 0.01), whereas the circularity of the spheroid was not affected by any tested compounds except SDS (which resulted in full disassembly of spheroid as mentioned above). No change in PI staining was observed for any treatment in St-T1b within the well-defined spheroid area, although local variations in PI intensity could be visually observed from the images for the cells protruding from the spheroid core (Figure 4). In the case of T-HESC spheroids, the compactness of the spheroid was significantly decreased in all treatments, as indicated by the apparent increase in the area occupied by the spheroid-forming cells (P < 0.001) and reduction of the spheroid circularity (P < 0.01) as compared to the negative control. Furthermore, the PI signal was significantly increased in T-HESC spheroids formed in the presence of either phthalate monoesters or SDS (P < 0.05), indicating compromised membrane integrity of the cells.

**Figure 3.**
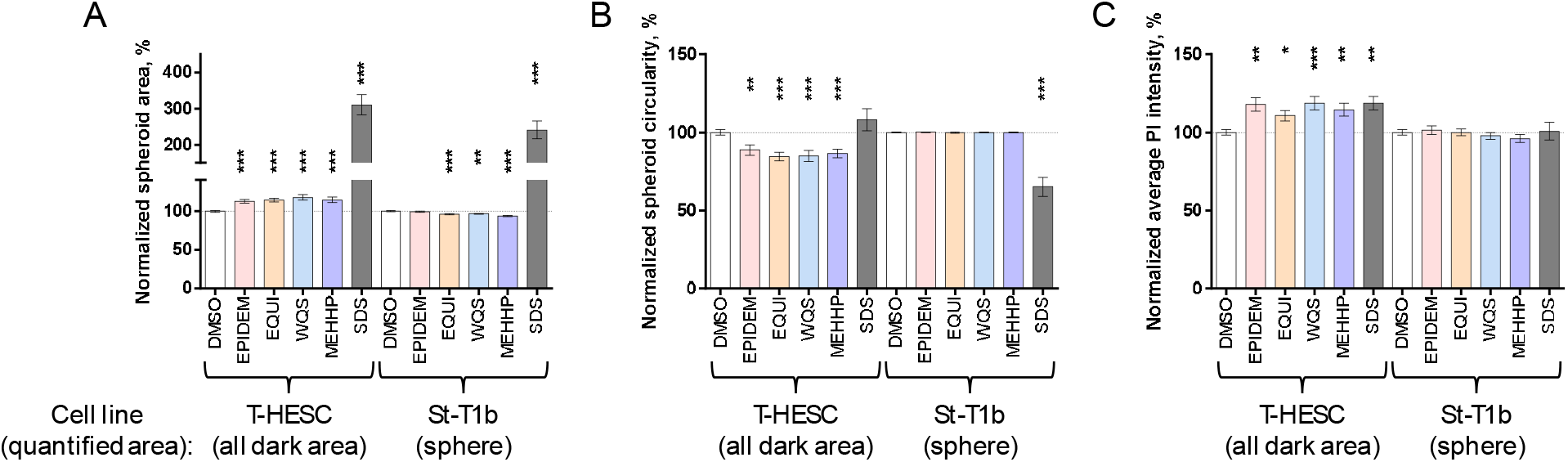
The presence of phthalate monoesters changes the phenotype of the spheroids formed from live T-HESC or St-T1b cells. A, normalized area; B, normalized circularity; C, normalized average intensity of PI staining. The average value of the indicated parameter ± SEM is shown (N ≥ 3 for each compound of interest in each cell line). The compounds of interest and the cell lines are listed below each graph. Net concentrations of phthalates: MEHHP – 21.6 μM, EPIDEM – 280 μM, EQUI – 330 μM, WQS – 220 μM. The dashed grey line indicates phenotype characteristic for the negative control (treatment with 0.1% DMSO). The asterisks indicate the statistical significance of difference between the phenotype in the absence *vs* presence of the compound of interest (only statistically significant comparisons are annotated); ^***^ indicates P < 0.001, ^**^ indicates P < 0.01, ^*^ indicates P < 0.05.

**Figure 4.**
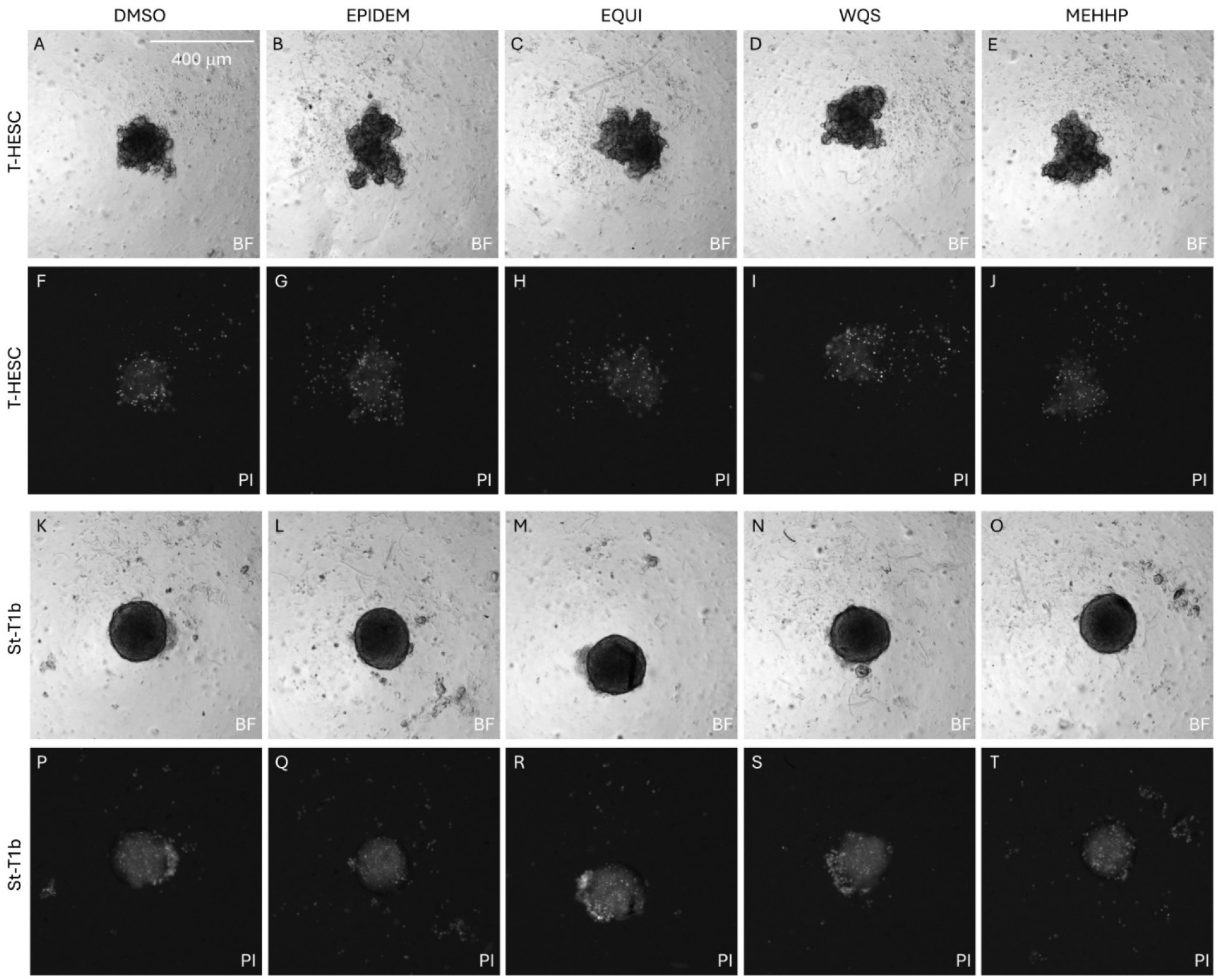
Imaging of spheroid formation in the presence of compounds of interest at 96 h after seeding of cells on to the ultra-low attachment plate. Data from a single representative experiment is shown. Cell lines are listed on the left: A-J, T-HESC; K-T, St-T1b. The compounds of interest are listed above the images: A, F, K and P, 0.1% DMSO (control); B, G, L and Q, EPIDEM; C, H, M and R, EQUI; D, I, N and S, WQS; E, J, O and T, MEHHP. Bright-field (BF) images are shown in A-E and K-O and propidium iodide staining (PI) in F-J and P-T. The scale bar is shown in the upper right corner in image A.

### 3.3. MEHHP interferes with tight junction assembly in T-HESC spheroids

Finally, we examined whether treatment with phthalate monoesters can alter the amount or localization pattern of characteristic tight junction proteins in spheroids. For that, we conducted 72-h spheroid formation as described above, followed by fixation and immunostaining; the nuclei were stained using Hoechst 33342. In a pilot experiment, we utilized both St-T1b and T-HESC cell lines and tried antibodies against three different tight junction proteins (39): claudin, occludin, and ZO-1. According to the imaging data presented in Supplementary Figure S3, ZO-1 yielded the highest signal in both cell lines and featured characteristic staining (40).

For the following experiments with phthalate monoester-treated spheroids, only T-HESC cell line was used as it yielded more statistically significant comparisons in the spheroid formation assay (Figure 3). In addition to the negative control (0.1% DMSO), as the compound of interest, only MEHHP was utilized, as the effects of all phthalate monoester-containing solutions were similar according to the spheroid formation assay (Figure 3). The characteristic microscopy images following 72-h treatment and immunostaining are presented in Figure 5A,B and Supplementary Figure S4, and the data on normalized ZO-1 signal intensity in 0.1% DMSO-*vs* MEHHP-treated spheroids is presented in Figure 5C (pooled data from three independent experiments). Overall, MEHHP treatment resulted in a significant reduction of ZO-1 signal (P < 0.01) and the formation of spotted rather than ridged ZO-1 localization pattern, thus confirming that phthalate monoesters can interfere with the formation of cell-cell contacts.

**Figure 5.**
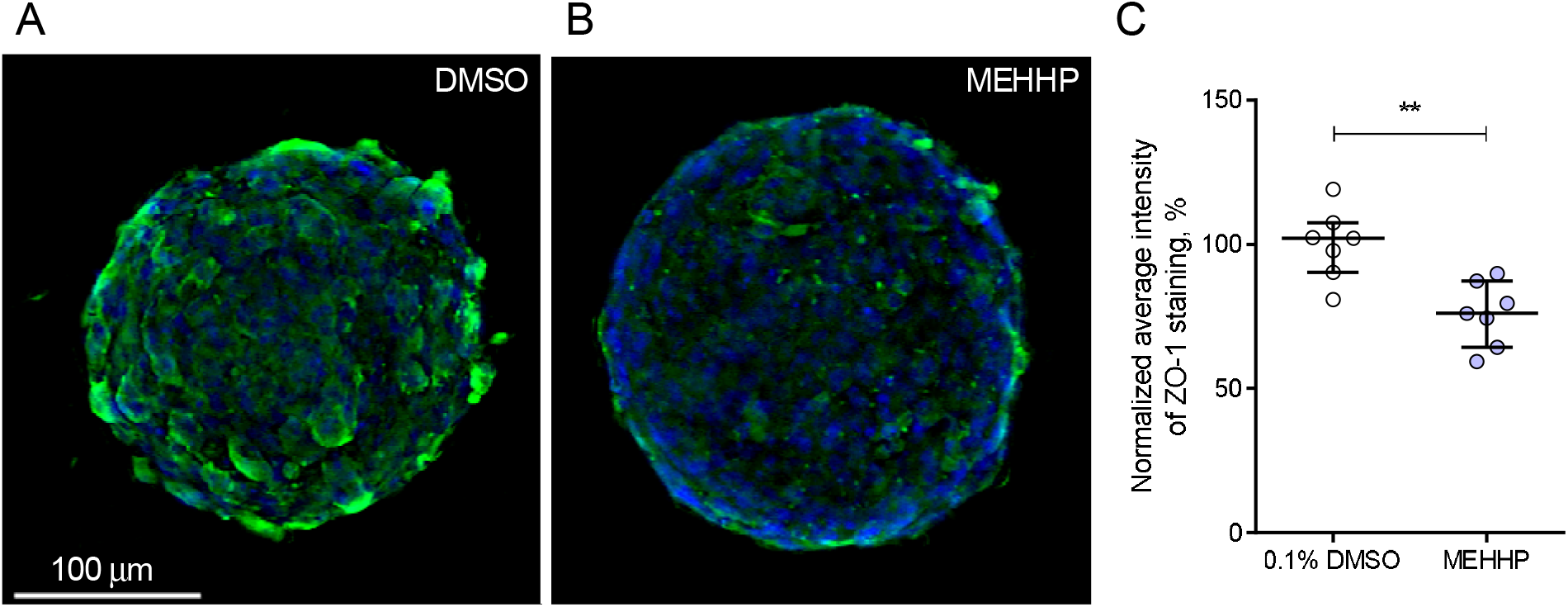
ZO-1 immunostaining in T-HESC spheroids reveals that MEHHP interferes with tight junction assembly. A and B, deconvolved fluorescence microscopy images from a single representative experiment showing the overlay of ZO-1 in the green channel and nuclear stain in the blue channel (10× objective). The compounds used for 72-h treatment are indicated in the top right (A: 0.1% DMSO, B: 21.6 μM MEHHP) and the scale bar is shown in the bottom left corner in A. C, quantification of ZO-1 average immunostaining intensity per spheroid area unit (pooled data, 2-3 spheroids per experiment, N = 3); circles denote individual spheroids, the middle line shows the median, and the whiskers protrude from the second to the third quartile. To compensate for the variation in staining conditions between the experiments, for each independent experiment, the ZO-1 signal was normalized to the mean signal observed in spheroids treated with 0.1% DMSO. The asterisks indicate the statistical significance of difference between the ZO-1 signal in the absence *vs* presence of MEHHP; ^**^ corresponds to P < 0.01.

## 4. Discussion

According to the Endocrine Society’s recent statement summarizing research on endocrine-disrupting chemicals performed during the years 2009-2014, phthalates are detectable in human urine, serum, and milk samples, and the estimated daily exposure to one major phthalate, di(2-ethylhexyl)phthalate (DEHP), ranges from 3 to 30 μg/kg (1). MEHHP is one of the major metabolites of DEHP, and MEHHP levels in urine or blood serum have been used as biomarkers for DEHP exposure assessment (41, 42).

The current study served as a follow-up to the previously published efforts characterizing the short-term effect of MEHHP and MEHHP-containing epidemiologically relevant mixtures on the transcriptome of cell lines and primary cells representing human endometrium (Visser *et al*., 2024, manuscript under revision), or human and mouse ovaries (11, 12). The latter reports indicated that phthalate monoesters are involved in multiple cellular pathways in both explored systems, with major differentially expressed genes related to cell adhesion, cytoskeleton and mitochondria in case of endometrium (Visser *et al*., 2024, manuscript under revision) and trans-membrane transport, metabolism, differentiation, chemotaxis, cell adhesion and kinase signalling in case of ovaries (11, 12). Here, our aim was to find the most general and intuitive common “denominator” explaining the versatility of MEHHP-affected cellular pathways observed by us and other research groups. We hypothesized that MEHHP can alter the fluidity of membranes in live cells – due to the structural similarity of MEHHP and some of the plasma membrane constituents and data from previous studies showing that endocrine disruptors can intercalate into the lipid bilayer (14–16). Alteration of membrane fluidity can, in turn, be propagated on the level of both cell-cell signalling and intracellular processes (20–25).

Using fluorescence anisotropy assay with probes that intercalate into bilayer at different depths and report on changes in membrane fluidity/rigidity, we found that even a short-term exposure of membranes prepared by lysis of T-HESC and St-T1b cells to MEHHP or MEHHP-containing mixtures is sufficient for statistically significant alteration in probe signal (Figure 1). Interestingly, different mixtures altered fluidity at different depths (Figure 2). This suggests that the presence of “shorter” phthalate monoesters (i.e., those incorporating substituents shorter than 8-carbon containing branched chain) in the incubation mixture modulates the MEHHP effect. This observation is highly relevant, given exposure of population to mixtures of endocrine disruptors rather than individual compounds. In the context of both reproduction-related and other models, phthalates and their metabolites have been previously reported to activate peroxisome proliferator-activated receptors (PPAR) (43–45). As the PPAR family is involved in cholesterol and triglyceride transport (46) and membrane fluidity regulation (47), the effect of phthalate monoesters on membrane fluidity can be augmented by interaction with PPAR – yet this is unlikely in the case of short-term incubations in simplified model systems, such as membrane preparations used in this study.

The evaluation of compounds of interest in a physiologically more relevant system represented by T-HESC and St-T1b spheroids indicated that the size and morphology of the forming spheroid were affected by the presence of phthalate monoesters. The outcomes of the spheroid formation assay were more dependent on the intrinsic tendency of the cell lines to form tight cell-cell contacts than on the treatment mixture (Figure 3): tightly packed spheroids tended to have smaller size upon 72-h treatment with phthalate monoesters, whereas loose spheroids tended to disintegrate. In our earlier research in the context of a very different model system (cancerous spheroids treated with a targeted drug), we have observed before that the readout of the assay is strongly dependent on the adhesion properties of the cell line used (30). The fact that MEHHP and various MEHHP-containing mixtures showed (within a single cell line) different impacts in the membrane fluidity study yet similar effect in the spheroid formation assay likely indicates that any kind of alteration in membrane fluidity perturbs the formation of cell-cell contacts. This notion and the reduction of ZO-1 immunostaining in MEHHP-treated spheroids confirmed in this work (Figure 5) are in line with the morphological changes reported in literature. Previous studies unveiled DEHP-induced increase of migratory properties and epithelial-mesenchymal transition (EMT) in human endometrial and endometriotic epithelial cells (48) and in breast cancer cell lines (49), whereas both increased migration and EMT are associated with loss of cell-cell contacts and drop in cellular adhesion. In a wider context, our findings might also explain the reported positive correlation between the phthalate exposure and endometriosis (50, 51), as the latter pathological condition is linked to the abnormal migratory properties of endometrial cells, as well as association of the phthalate exposure with adverse IVF outcomes (52, 53), as cell-cell adhesion is crucial for embryo implantation.

The current study has several limitations, including the use of a single concentration of compounds of interest and exploring the effects of compounds in cell lines instead of primary cells. These methodological choices were made due to the characteristics of the membrane fluidity assay, which requires a large amount of biological material to achieve a sufficient measurement window. Different assays also required different exposure times, as biological stability and homogeneity of the membrane preparations does not enable long incubations, while spheroids are formed on the scale of days rather than hours. Furthermore, the conclusions regarding interference of MEHHP with tight junction assembly were made based on immunostaining of a single protein in a single cell line. The reason for such experimental design was that immunostaining of spheroids is technically substantially more demanding than immunostaining of adherent cultures, and a limited set of parameters had to be chosen for performing a sufficient number of independent experiments. Still, we demonstrated successfully that phthalate monoesters can affect membrane fluidity and cell-cell contacts, which explains versatile downstream effects uncovered in a multitude of previous studies. Given the recent advances in the techniques suitable for the examination of lipid ensembles, functions, and trafficking, more attention must be paid to the role of endocrine disruptors as chemicals that can affect lipidome.

## Acknowledgements

We express our deepest gratitude to Dr. Birgit Gellersen and Dr. Martin Götte for providing the St-T1b cell line, and to Dr. Santa Veikšina for valuable advice regarding the preparation of membrane homogenates.

## Author contributions

D.L. formulated the initial hypothesis, performed all membrane fluidity and spheroid formation experiments together with the corresponding data analysis, and drafted the manuscript; K.K. performed assembly and immunostaining of spheroids; S.K. participated in setting up of membrane fluidity assay, performed spheroid imaging and imaging data deconvolution; N.V. participated in the assessment of phthalate mixture quality and establishment of cell culturing conditions; M.L. and J.A.F. prepared and validated the phthalate mixtures; T.K.K., M.O. and P.D. participated in generation of study concept, acquired the necessary resources, and supervised the PhD student N.V.; A.S. was responsible for project administration, participated in the generation of study concept, and acquired the necessary resources. All authors read and approved the final manuscript.

## References

1. Gore, A. C., Chappell, V. A., Fenton, S. E., Flaws, J. A., Nadal, A., Prins, G. S., Toppari, J., and Zoeller, R. T. (2015) EDC-2: The Endocrine Society’s Second Scientific Statement on Endocrine-Disrupting Chemicals. Endocr Rev 36, E1–E150

2. Tan, H., Chen, Q., Hong, H., Benfenati, E., Gini, G. C., Zhang, X., Yu, H., and Shi, W. (2021) Structures of Endocrine-Disrupting Chemicals Correlate with the Activation of 12 Classic Nuclear Receptors. Environ. Sci. Technol. 55, 16552–16562

3. An, J., Roh, H.-H., Jeong, H., Lee, K.-Y., and Rhim, T. (2023) Rapid Assessment of Di(2-ethylhexyl) Phthalate Migration from Consumer PVC Products. Toxics 12, 7

4. Jang, E. A., Kim, K. N., and Bae, S. H. (2024) Associations of concentrations of eight urinary phthalate metabolites with the frequency of use of common adult consumer and personal-care products. Sci Rep 14, 5187

5. Wang, Z., Ma, J., Wang, T., Qin, C., Hu, X., Mosa, A., and Ling, W. (2023) Environmental health risks induced by interaction between phthalic acid esters (PAEs) and biological macromolecules: A review. Chemosphere 328, 138578

6. Zhou, C., Gao, L., and Flaws, J. A. (2017) Exposure to an Environmentally Relevant Phthalate Mixture Causes Transgenerational Effects on Female Reproduction in Mice. Endocrinology 158, 1739–1754

7. Arrigo, F., Impellitteri, F., Piccione, G., and Faggio, C. (2023) Phthalates and their effects on human health: Focus on erythrocytes and the reproductive system. Comparative Biochemistry and Physiology Part C: Toxicology & Pharmacology 270, 109645

8. Mankidy, R., Wiseman, S., Ma, H., and Giesy, J. P. (2013) Biological impact of phthalates. Toxicology Letters 217, 50–58

9. Wang, Y. and Qian, H. (2021) Phthalates and Their Impacts on Human Health. Healthcare 9, 603

10. Scholz, N. (2003) Ecotoxicity and biodegradation of phthalate monoesters. Chemosphere 53, 921–926

11. Tarvainen, I., Soto, D. A., Laws, M. J., Björvang, R. D., Damdimopoulos, A., Roos, K., Li, T., Kramer, S., Li, Z., Lavogina, D., Visser, N., Kallak, T. K., Lager, S., Gidlöf, S., Edlund, E., Papaikonomou, K., Öberg, M., Olovsson, M., Salumets, A., Velthut-Meikas, A., Flaws, J. A., and Damdimopoulou, P. (2023) Identification of phthalate mixture exposure targets in the human and mouse ovary in vitro. Reprod Toxicol 119, 108393

12. Panagiotou, E. M., Damdimopoulos, A., Li, T., Moussaud-Lamodière, E., Pedersen, M., Lebre, F., Pettersson, K., Arnelo, C., Papaikonomou, K., Alfaro-Moreno, E., Lindskog, C., Svingen, T., and Damdimopoulou, P. (2024) Exposure to the phthalate metabolite MEHP impacts survival and growth of human ovarian follicles in vitro. Toxicology 505, 153815

13. Lavogina, D., Stepanjuk, A., Peters, M., Samuel, K., Kasvandik, S., Khatun, M., Arffman, R. K., Enkvist, E., Viht, K., Kopanchuk, S., Lättekivi, F., Velthut-Meikas, A., Uri, A., Piltonen, T. T., Rinken, A., and Salumets, A. (2021) Progesterone triggers Rho kinase-cofilin axis during in vitro and in vivo endometrial decidualization. Hum Reprod 36, 2230–2248

14. Kwon, J.-H., Liljestrand, H. M., and Katz, L. E. (2006) Partitioning of moderately hydrophobic endocrine disruptors between water and synthetic membrane vesicles. Environ Toxicol Chem 25, 1984–1992

15. Horváth, Á., Erostyák, J., and Szőke, É. (2022) Effect of Lipid Raft Disruptors on Cell Membrane Fluidity Studied by Fluorescence Spectroscopy. Int J Mol Sci 23, 13729

16. Katata, V. M., Maximino, M. D., Silva, C. Y., and Alessio, P. (2022) The Role of Cholesterol in the Interaction of the Lipid Monolayer with the Endocrine Disruptor Bisphenol-A. Membranes 12, 729

17. Okamura, E., Wakai, C., Matubayasi, N., Sugiura, Y., and Nakahara, M. (2004) Limited slowdown of endocrine-disruptor diffusion in confined fluid lipid membranes. Phys Rev Lett 93, 248101

18. Izbicka, E. and Streeper, R. T. (2021) Adaptive Membrane Fluidity Modulation: A Feedback Regulated Homeostatic System Hiding in Plain Sight. In Vivo 35, 2991–3000

19. Louis, M., Tahrioui, A., Verdon, J., David, A., Rodrigues, S., Barreau, M., Manac’h, M., Thiroux, A., Luton, B., Dupont, C., Calvé, M. L., Bazire, A., Crépin, A., Clabaut, M., Portier, E., Taupin, L., Defontaine, F., Clamens, T., Bouffartigues, E., Cornelis, P., Feuilloley, M., Caillon, J., Dufour, A., Berjeaud, J.-M., Lesouhaitier, O., and Chevalier, S. (2022) Effect of Phthalates and Their Substitutes on the Physiology of Pseudomonas aeruginosa. Microorganisms 10, 1788

20. Sunshine, H. and Iruela-Arispe, M. L. (2017) Membrane Lipids and Cell Signaling. Curr Opin Lipidol 28, 408–413

21. Deng, B., Shen, W.-J., Dong, D., Azhar, S., and Kraemer, F. B. (2019) Plasma membrane cholesterol trafficking in steroidogenesis. FASEB J 33, 1389–1400

22. Ray, S., Kassan, A., Busija, A. R., Rangamani, P., and Patel, H. H. (2016) The plasma membrane as a capacitor for energy and metabolism. Am J Physiol Cell Physiol 310, C181– C192

23. McCusker, D. and Kellogg, D. R. (2012) Plasma membrane growth during the cell cycle: unsolved mysteries and recent progress. Curr Opin Cell Biol 24, 845–851

24. Jena, B. P. (2007) Secretion machinery at the cell plasma membrane. Curr Opin Struct Biol 17, 437–443

25. Prinz, W. A., Toulmay, A., and Balla, T. (2020) The functional universe of membrane contact sites. Nat Rev Mol Cell Biol 21, 7–24

26. Samalecos, A., Reimann, K., Wittmann, S., Schulte, H. M., Brosens, J. J., Bamberger, A.-M., and Gellersen, B. (2009) Characterization of a novel telomerase-immortalized human endometrial stromal cell line, St-T1b. Reprod Biol Endocrinol 7, 76

27. Kopanchuk, S., Vavers, E., Veiksina, S., Ligi, K., Zvejniece, L., Dambrova, M., and Rinken, A. (2022) Intracellular dynamics of the Sigma-1 receptor observed with super-resolution imaging microscopy. PLOS ONE 17, e0268563

28. Kopanchuk, S. and Rinken, A. (2001) Changes in Membrane Fluidity During the Micelle Formation Determine the Efficiency of the Solubilization of Muscarinic Receptors. Proceedings of the Estonian Academy of Sciences 50, 229–240

29. Schindelin, J., Arganda-Carreras, I., Frise, E., Kaynig, V., Longair, M., Pietzsch, T., Preibisch, S., Rueden, C., Saalfeld, S., Schmid, B., Tinevez, J.-Y., White, D. J., Hartenstein, V., Eliceiri, K., Tomancak, P., and Cardona, A. (2012) Fiji: an open-source platform for biological-image analysis. Nat Methods 9, 676–682

30. Lavogina, D., Krõlov, M. K., Vellama, H., Modhukur, V., Di Nisio, V., Lust, H., Eskla, K.-L., Salumets, A., and Jaal, J. (2024) Inhibition of epigenetic and cell cycle-related targets in glioblastoma cell lines reveals that onametostat reduces proliferation and viability in both normoxic and hypoxic conditions. Sci Rep 14, 4303

31. Soulez, F., Denis, L., Tourneur, Y., and Thiébaut, É. (2012) Blind deconvolution of 3D data in wide field fluorescence microscopy. presented at the 2012 9th IEEE International Symposium on Biomedical Imaging (ISBI)

32. de Chaumont, F., Dallongeville, S., Chenouard, N., Hervé, N., Pop, S., Provoost, T., Meas-Yedid, V., Pankajakshan, P., Lecomte, T., Le Montagner, Y., Lagache, T., Dufour, A., and Olivo-Marin, J.-C. (2012) Icy: an open bioimage informatics platform for extended reproducible research. Nat Methods 9, 690–696

33. Nelson, S. C., Neeley, S. K., Melonakos, E. D., Bell, J. D., and Busath, D. D. (2012) Fluorescence anisotropy of diphenylhexatriene and its cationic Trimethylamino derivative in liquid dipalmitoylphosphatidylcholine liposomes: opposing responses to isoflurane. BMC Biophys 5, 5

34. Marczak, A. (2009) Fluorescence anisotropy of membrane fluidity probes in human erythrocytes incubated with anthracyclines and glutaraldehyde. Bioelectrochemistry 74, 236–239

35. do Canto, A. M. T. M., Robalo, J. R., Santos, P. D., Carvalho, A. J. P., Ramalho, J. P. P., and Loura, L. M. S. (2016) Diphenylhexatriene membrane probes DPH and TMA-DPH: A comparative molecular dynamics simulation study. Biochimica et Biophysica Acta (BBA) - Biomembranes 1858, 2647–2661

36. Neves, P., Lopes, S. C. D. N., Sousa, I., Garcia, S., Eaton, P., and Gameiro, P. (2009) Characterization of membrane protein reconstitution in LUVs of different lipid composition by fluorescence anisotropy. Journal of Pharmaceutical and Biomedical Analysis 49, 276–281

37. Wong, C.-W., Han, H.-W., and Hsu, S. (2022) Changes of cell membrane fluidity for mesenchymal stem cell spheroids on biomaterial surfaces. World Journal of Stem Cells 14, 616

38. Wu, H., Yu, M., Miao, Y., He, S., Dai, Z., Song, W., Liu, Y., Song, S., Ahmad, E., Wang, D., and Gan, Y. (2019) Cholesterol-tuned liposomal membrane rigidity directs tumor penetration and anti-tumor effect. Acta Pharmaceutica Sinica B 9, 858–870

39. Kuo, W.-T., Odenwald, M. A., Turner, J. R., and Zuo, L. (2022) Tight junction proteins occludin and ZO-1 as regulators of epithelial proliferation and survival. Ann N Y Acad Sci 1514, 21–33

40. Troitskaya, O., Novak, D., Nushtaeva, A., Savinkova, M., Varlamov, M., Ermakov, M., Richter, V., and Koval, O. (2021) EGFR Transgene Stimulates Spontaneous Formation of MCF7 Breast Cancer Cells Spheroids with Partly Loss of HER3 Receptor. International Journal of Molecular Sciences 22, 12937

41. Silva, M. J., Reidy, J. A., Preau, J. L., Samandar, E., Needham, L. L., and Calafat, A. M. (2006) Measurement of eight urinary metabolites of di(2-ethylhexyl) phthalate as biomarkers for human exposure assessment. Biomarkers 11, 1–13

42. Kato, K., Silva, M. J., Reidy, J. A., Hurtz, D., Malek, N. A., Needham, L. L., Nakazawa, H., Barr, D. B., and Calafat, A. M. (2004) Mono(2-ethyl-5-hydroxyhexyl) phthalate and mono-(2-ethyl-5-oxohexyl) phthalate as biomarkers for human exposure assessment to di-(2-ethylhexyl) phthalate. Environ Health Perspect 112, 327–330

43. Hurst, C. H. and Waxman, D. J. (2003) Activation of PPARalpha and PPARgamma by environmental phthalate monoesters. Toxicol Sci 74, 297–308

44. Lapinskas, P. J., Brown, S., Leesnitzer, L. M., Blanchard, S., Swanson, C., Cattley, R. C., and Corton, J. C. (2005) Role of PPARalpha in mediating the effects of phthalates and metabolites in the liver. Toxicology 207, 149–163

45. Meling, D. D., De La Torre, K. M., Arango, A. S., Gonsioroski, A., Deviney, A. R. K., Neff, A. M., Laws, M. J., Warner, G. R., Tajkhorshid, E., and Flaws, J. A. (2022) Phthalate monoesters act through peroxisome proliferator-activated receptors in the mouse ovary. Reproductive Toxicology 110, 113–123

46. Barbier, O., Torra, I. P., Duguay, Y., Blanquart, C., Fruchart, J.-C., Glineur, C., and Staels, B. (2002) Pleiotropic actions of peroxisome proliferator-activated receptors in lipid metabolism and atherosclerosis. Arterioscler Thromb Vasc Biol 22, 717–726

47. Wu, Y., Tan, X., Tian, J., Liu, X., Wang, Y., Zhao, H., Yan, Z., Liu, H., and Ma, X. (2017) PPARγ Agonist Ameliorates the Impaired Fluidity of the Myocardial Cell Membrane and Cardiac Injury in Hypercholesterolemic Rats. Cardiovasc Toxicol 17, 25–34

48. Kim, H. G., Lim, Y. S., Hwang, S., Kim, H.-Y., Moon, Y., Song, Y. J., Na, Y.-J., and Yoon, S. (2022) Di-(2-ethylhexyl) Phthalate Triggers Proliferation, Migration, Stemness, and Epithelial–Mesenchymal Transition in Human Endometrial and Endometriotic Epithelial Cells via the Transforming Growth Factor-β/Smad Signaling Pathway. International Journal of Molecular Sciences 23, 3938

49. Khan, N. G., Eswaran, S., Adiga, D., Sriharikrishnaa, S., Chakrabarty, S., Rai, P. S., and Kabekkodu, S. P. (2022) Integrated bioinformatic analysis to understand the association between phthalate exposure and breast cancer progression. Toxicology and Applied Pharmacology 457, 116296

50. Kim, J. H. and Kim, S. H. (2020) Exposure to Phthalate Esters and the Risk of Endometriosis. Dev Reprod 24, 71–78

51. Chou, Y.-C. and Tzeng, C.-R. (2021) The impact of phthalate on reproductive function in women with endometriosis. Reproductive Medicine and Biology 20, 159–168

52. Al-Saleh, I., Coskun, S., Al-Doush, I., Abduljabbar, M., Al-Rouqi, R., Al-Rajudi, T., and Al-Hassan, S. (2019) Couples exposure to phthalates and its influence on in vitro fertilization outcomes. Chemosphere 226, 597–606

53. Begum, T. F., Fujimoto, V. Y., Gerona, R., McGough, A., Lenhart, N., Wong, R., Mok-Lin, E., Melamed, J., Butts, C. D., and Bloom, M. S. (2021) A pilot investigation of couple-level phthalates exposure and in vitro fertilization (IVF) outcomes. Reprod Toxicol 99, 56–64

