## Supplementary material for "Phthalate monoesters affect membrane fluidity and cell-cell contacts in endometrial stromal cell lines": Table S1

***Supplementary data***

Table S1. Composition of phthalate monoester stock solutions

| Stock: | EPIDEM, theoretical | | | EPIDEM | EQUI, theoretical | | | EQUI | WQS, theoretical | | | WQS | MEHHP, theoretical | | |
| --- | --- | --- | --- | --- | --- | --- | --- | --- | --- | --- | --- | --- | --- | --- | --- |
|  | % ^a^ | mg/mL | mM | mM ^b^ | % ^a^ | mg/mL | mM | mM ^b^ | % ^a^ | mg/mL | mM | mM ^b^ | % ^a^ | mg/mL | mM |
| MBP | 6.7 | 3.97 | 17.9 | 13 ± 3 | 11.1 | 6.59 | 29.7 | 21 ± 5 | - | - | - | - | - | - | - |
| MBzP | 2.6 | 1.78 | 6.94 | 14 ± 3 | 11.1 | 7.59 | 29.7 | 50 ± 5 | 3.2 | 1.78 | 6.94 | 13 ± 2 | - | - | - |
| MCPP | 0.8 | 0.54 | 2.14 | 1.6 ± 0.2 | 11.1 | 7.48 | 29.7 | 26 ± 8 | - | - | - | - | - | - | - |
| MECPP | 6.4 | 5.26 | 17.1 | 19 ± 4 | 11.1 | 9.13 | 29.7 | 33 ± 3 | - | - | - | - | - | - | - |
| MEHHP | 8.1 | 6.36 | 21.6 | 19 ± 5 | 11.1 | 8.72 | 29.7 | 26 ± 3 | 9.9 | 6.36 | 21.6 | 18 ± 2 | 100 | 6.36 | 21.6 |
| MEHP | 1.3 | 0.97 | 3.47 | 7.0 ± 1.2 | 11.1 | 8.24 | 29.7 | 44 ± 5 | - | - | - | - | - | - | - |
| MEOHP | 3.1 | 2.42 | 8.28 | 16 ± 3 | 11.1 | 8.66 | 29.7 | 58 ± 5 | - | - | - | - | - | - | - |
| MEP | 65.6 | 34.0 | 175 | 170 ± 30 | 11.1 | 5.76 | 29.7 | 34 ± 6 | 80.3 | 34.0 | 175 | 170 ± 10 | - | - | - |
| MiBP | 5.3 | 3.14 | 14.2 | 21 ± 6 | 11.1 | 6.59 | 29.7 | 42 ± 6 | 6.5 | 3.14 | 14.2 | 21 ± 3 | - | - | - |
| SUM | **100** | **58.4** | **267** ^c^ | **280** ^d^ | **100** | **68.8** | **267** ^c^ | **330** ^d^ | **100** | **45.3** | **218** ^c^ | **220** ^d^ | **100** | **6.36** | **21.6** ^c^ |

^a^ Theoretical molar percent of an individual compound from all phthalate monoesters contained in the stock solution; ^b^ calculated average concentration of an individual compound from all phthalate monoesters contained in the stock solution based on data measured experimentally for spent culture media in previous studies (PMID 37160244 and Visser *et al*., 2024, manuscript under revision); ^c^ theoretical total molar concentration of all dissolved compounds in the stock solution; ^d^ calculated total molar concentration of all dissolved compounds in the stock solution based on experimental data. Abbreviations: EQUI, equimolar mixture; EPIDEM, epidemiological mixture; MBP, monobutyl phthalate; MBzP, monobenzyl phthalate; MCPP, mono(3-carboxypropyl) phthalate; MECPP, mono(5-carboxy-2ethylpentyl) phthalate; MEHHP, mono(2-ethyl-5-hydroxyhexyl) phthalate; MEHP, mono(ethylhexyl) phthalate; MEOHP, mono(2-ethyl-5-oxyhexyl) phthalate; MEP, monoethyl phthalate; MiBP, monoisobutyl phthalate; WQS, weighted quantile sum-based mixture; - means that the compound is absent from the solution. All stock solutions were used at 1:1000 final dilution in this study.
