## Supplementary material for "Phthalate monoesters affect membrane fluidity and cell-cell contacts in endometrial stromal cell lines": Figure S1

***Supplementary data***

Figure S1. Membrane fluidity measurement at the presence of SDS dilution series

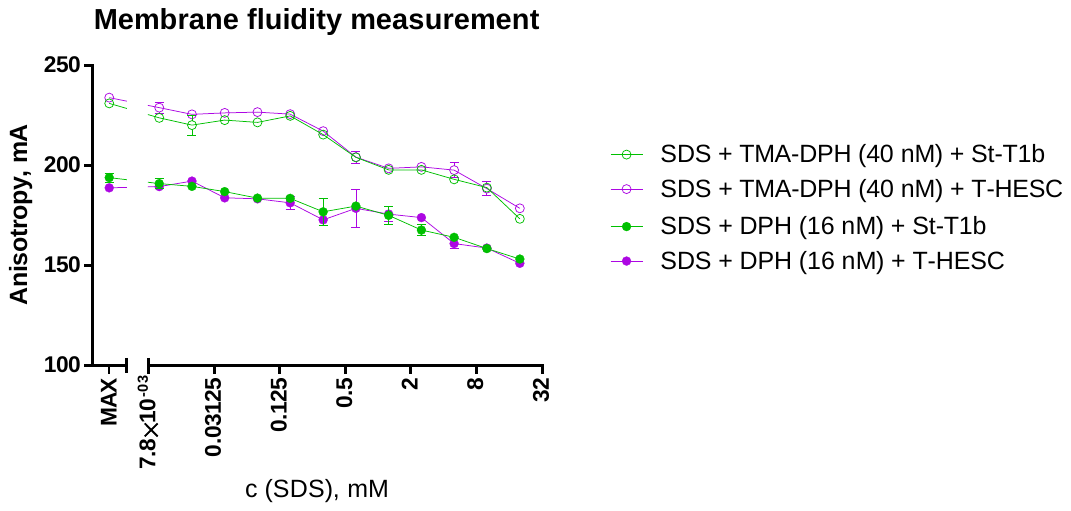

Data from a single pilot experiment is shown (average anisotropy value ± SEM for duplicates). Full circles show data for the DPH probe and open circles for the TMA-DPH probe; different colours indicate membrane preparations obtained from St-T1b cells (green) or T-HESC cells (magenta). MAX indicates the sample containing probe and membranes but no SDS. Please note that the y-axis does not start from zero.
