## Supplementary material for "Phthalate monoesters affect membrane fluidity and cell-cell contacts in endometrial stromal cell lines": Figure S2

***Supplementary data***

Figure S2. Imaging of spheroid formation at various time points after seeding of cells onto the ultra-low attachment plate

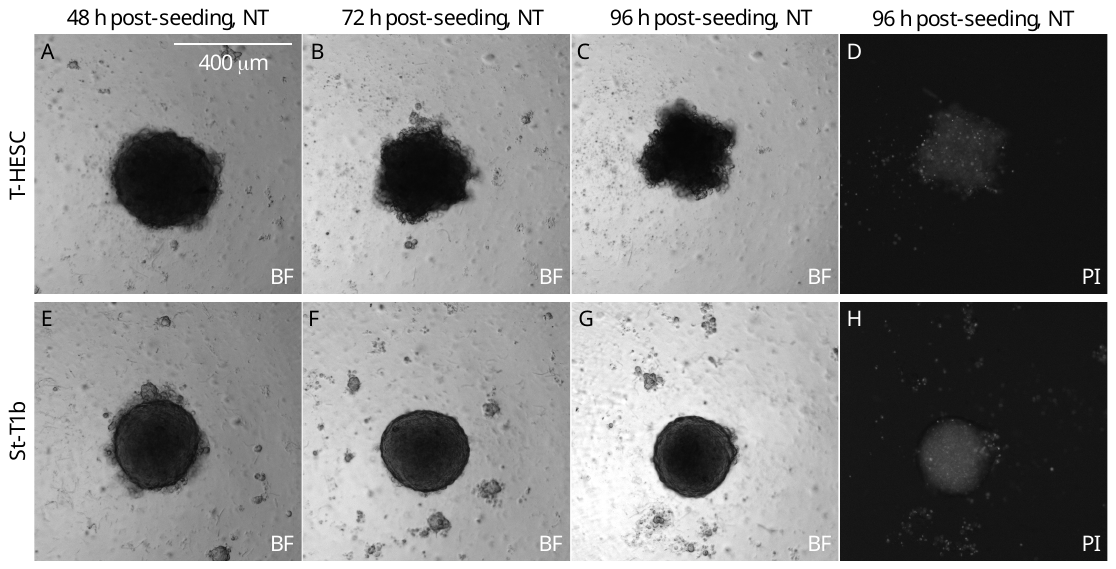

Data from a single pilot experiment is shown (non-treated spheroids). Cell lines are listed on the left: A-D, T-HESC; E-H, St-T1b. Time points post-seeding are listed above the images: A and E, 48 h; B and F, 72 h; C-D and G-H, 96 h. Bright-field (BF) images are shown in A-C and E-G and propidium iodide staining (PI) in D and H. The scale bar is shown in the upper right corner in image A.
