## Supplementary material for "Phthalate monoesters affect membrane fluidity and cell-cell contacts in endometrial stromal cell lines": Figure S3

***Supplementary data***

Figure S3. Immunostaining of tight junction proteins in fixed spheroids

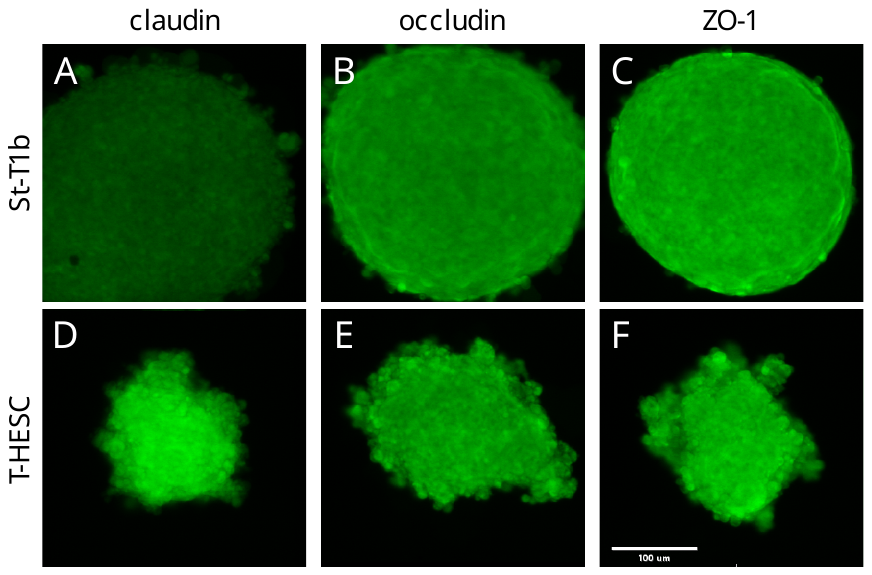

Microscopy data from a single pilot experiment is shown (non-treated spheroids, non-deconvolved images, 10× objective). Cell lines: A-C, St-T1b; D-F, T-HESC. Immunostaining: A and D, claudin; B and E, occludin; C and F, ZO-1. The scale bar is shown in the bottom right in image F.
