## Supplementary material for "Phthalate monoesters affect membrane fluidity and cell-cell contacts in endometrial stromal cell lines": Figure S4

***Supplementary data***

Figure S4. Immunostaining of ZO-1 and nuclear staining in fixed T-HESC spheroids

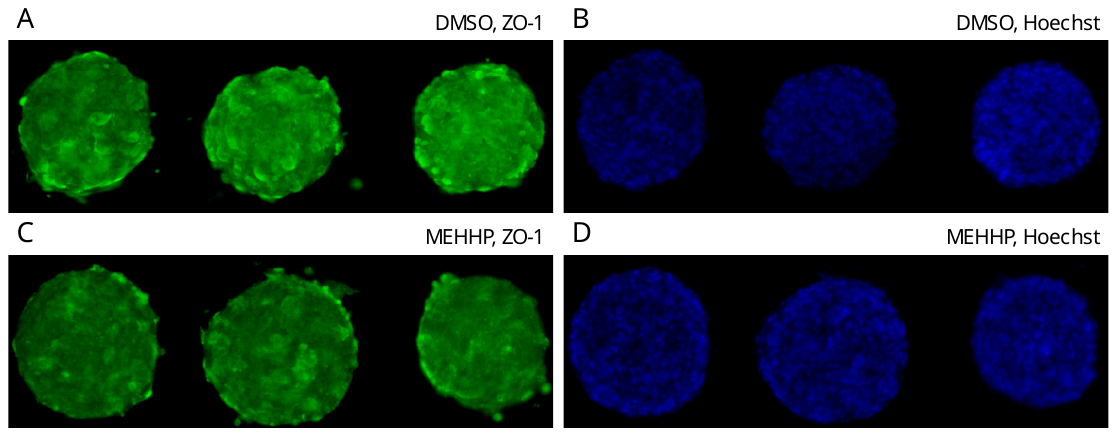

Microscopy data from a single representative experiment is shown (non-deconvolved images, 10× objective, individual channels). Each panel features staining for three replicate spheroids. Treatment (72 h): A and B, 0.1% DMSO; C and D, MEHHP. Channels: A and C, ZO-1; B and D, Hoechst 33342.
